# IMiQC: a novel protein quality control compartment protecting mitochondrial functional integrity

**DOI:** 10.1101/075127

**Authors:** Michael Bruderek, Witold Jaworek, Anne Wilkening, Cornelia Rüb, Giovanna Cenini, Marc Sylvester, Wolfgang Voos

## Abstract

Aggregation processes can cause severe perturbations of cellular homeostasis and are frequently associated with diseases. We performed a comprehensive analysis of mitochondrial quality and function in presence of misfolded, aggregation-prone polypeptides. Although we observed significant aggregate formation inside mitochondria, we observed only a minor impairment of mitochondrial function. We could show that detoxification of misfolded reporter polypeptides as well as endogenous proteins inside mitochondria takes place via their sequestration into the specific organellar deposit site <u>I</u>ntra-<u>M</u>itochondrial Protein <u>Q</u>uality Control <u>C</u>ompartment (IMiQC). Only minor amounts of co-aggregated proteins were associated with IMiQC and neither resolubilization nor degradation by the mitochondrial PQC system were observed. The single IMiQC aggregate deposit was not transferred to daughter cells during cell division. Detoxification of misfolded polypeptides via IMiQC formation was highly dependent on a functional mitochondrial fission machinery. We conclude that the formation of the aggregate deposit is an important mechanism to maintain full functionality of mitochondria under proteotoxic stress conditions.

List of Abbreviations

2D-Page: 2-dimensional polyacrylamide gel electrophoresis;
Δψ_mt_: electric potential across the inner mitochondrial membrane;
AA: amino acid;
BN-PAGE: blue-native polyacrylamide gel electrophoresis;
DHFR: dihydrofolate reductase;
GFP: green fluorescent protein;
IMM: inner mitochondrial membrane;
IMS: intermembrane space;
MPP: mitochondrial processing peptidase;
MTS: mitochondrial targeting sequence;
OMM: outer mitochondrial membrane;
SDm: standard error of the mean;
SDS-PAGE: sodiumdodecylsulfate polyacrylamide gel electrophoresis;
TCA: trichloroacetic acid;
TOM: translocase of the outer membrane;
TIM: translocase of the inner membrane;
TMRE: tetramethylrhodamine ethyl ester.

## Introduction

Cellular survival depends on the maintenance of protein function under normal and stress conditions, a process summarized as protein homeostasis. Recent evidence indicated that a formation of large aggregate deposits might represent a protective mechanism to counteract proteotoxic stress conditions ^1^. An organized formation of aggregate deposits seems to be a regulated, cell-protective process that sequesters potential toxic molecules from the rest of the cellular environment and also may allow a specific aggregate removal ^2^. In bacteria, large amounts of misfolded proteins, in particular upon overexpression, accumulate in form of one or two relatively large inclusion bodies that are typically localized at the cell poles ^3^. In the yeast *Saccharomyces cerevisiae*, three different classes of aggregate deposits have been identified so far ^4^. Upon proteotoxic stress, cytosolic misfolded proteins are either transported into the nucleus, forming INQ (intranuclear quality control compartment) ^5, 6^, or remain in the cytosol and are arranged as CytoQ compartments. CytoQ represent peripheral stress-induced aggregate sites ^7^, also termed ‘stress foci’ ^8^ or ‘Q-bodies’ ^9^ Aggregated proteins located in CytoQ or INQ may be resolubilized by the cytosolic disaggregase machinery consisting of members of the Hsp70 and Hsp100 chaperone families. In contrast, amyloid-like aggregates, formed by β-sheet rich proteins, are stored separately in a compartment localized next to the vacuole, called IPOD (insoluble protein deposit) ^5^. Deposition sites for protein aggregates have also been described in mammalian cells. Overexpressed, folding-deficient proteins accumulate in the aggresome located in proximity of the microtuble-organizing centre (MTOC) ^10^. The deposition of aggregated polypeptides at specific sites inside a cell may fulfill at least three different purposes. i) The misfolded polypeptides are sequestered from the cellular environment and thereby neutralized. ii) The localization of the deposits facilitates an eventual removal of the aggregates. iii) An asymmetric distribution of damaged polypeptides at few or a single site inside the cell might prevent the transfer of the potentially toxic molecules to the daughter cell during cell division. Generally, an asymmetric distribution of aging factors seems to be an important aspect of cellular rejuvenation and determination of life expectancy ^11^.

Due to the endosymbiotic origin of mitochondria, protein homeostasis on the polypeptide level is maintained by an endogenous protein-quality control (PQC) system, composed of molecular chaperones and ATP-dependent proteases ^12^. In addition, cells exhibit additional mechanisms acting on the organellar level to maintain mitochondrial quality. In case of an accumulation of misfolded polypeptides in mitochondria the mitochondrial unfolded proteins response (UPR^mt^) may induce the increased expression of mitochondrial PQC components ^13^. Cells also have the ability to specifically remove dysfunctional mitochondria as a whole by a variation of autophagy, termed mitophagy ^14^. Mitophagy reactions are closely connected with another important cellular mechanism of mitochondrial quality control, the potential of mitochondria to undergo fission and fusion reactions. In addition to the involvement in mitophagy, mitochondrial dynamics contribute to a complementation of mtDNA mutations ^15^.

In this work, we expressed a folding-deficient, aggregation-prone reporter protein in mitochondria of the *Saccharomyces cerevisiae* to characterize mitochondria-specific proteotoxic effects. We analyzed the aggregation behavior of the fluorescent reporter by biochemical assays as well as by a microscopic analysis of mitochondrial morphology and distribution. We show that aggregation-prone mitochondrial polypeptides are sequestered into a large aggregate deposit site, thereby preventing overall proteotoxic damage to mitochondrial functions. We characterized the effects of mitochondrial fusion and fission components on aggregate deposit formation and analyzed transfer of aggregates to daughter cells during cell division. We conclude that aggregate deposition represents a secondary protection mechanism under proteotoxic stress, when the reactivity of the mitochondrial PQC system as primary system is overwhelmed.

## Results

### Destabilized proteins accumulate in the <u>I</u>ntra-<u>M</u>itochondrial Protein <u>Q</u>uality Compartment (IMiQC)

To test the sensitivity of mitochondria to the expression of aggregation-prone proteins, we designed a fusion protein, termed *mt*GFP-DHFR_ds_, which consisted of GFP and the dihydrofolate reductase (DHFR) from mouse, containing three specific mutations (Cys7Ser; Ser42Cys; Arg49Cys) at sites required for protein stability. The fusion protein was targeted to the mitochondrial matrix due to an N-terminal mitochondrial targeting sequence (MTS) derived from the yeast mitochondrial protein cytochrome *b_2_* (Fig. 1A). As controls, we used two mitochondrially-targeted fusion proteins containing either GFP fused to non-mutated DHFR or GFP alone with an N-terminal MTS (*mt*GFP-DHFR and *mt*GFP). The expression of all GFP-fusion proteins was under the control of a galactose-inducible promoter (Fig. 1A; Sup. 1A). After expression, all reporter proteins were resistant against externally added proteinase K (PK), similar to the endogenous mitochondrial protein Tim23, indicating their localization inside mitochondria (Fig. 1B). As control, the mitochondrial outer membrane protein Tom40 was digested by PK treatment. After detergent lysis of the mitochondria, both the reporter constructs as well as Tim23 became sensitive to protease digestion.

**Figure 1.**
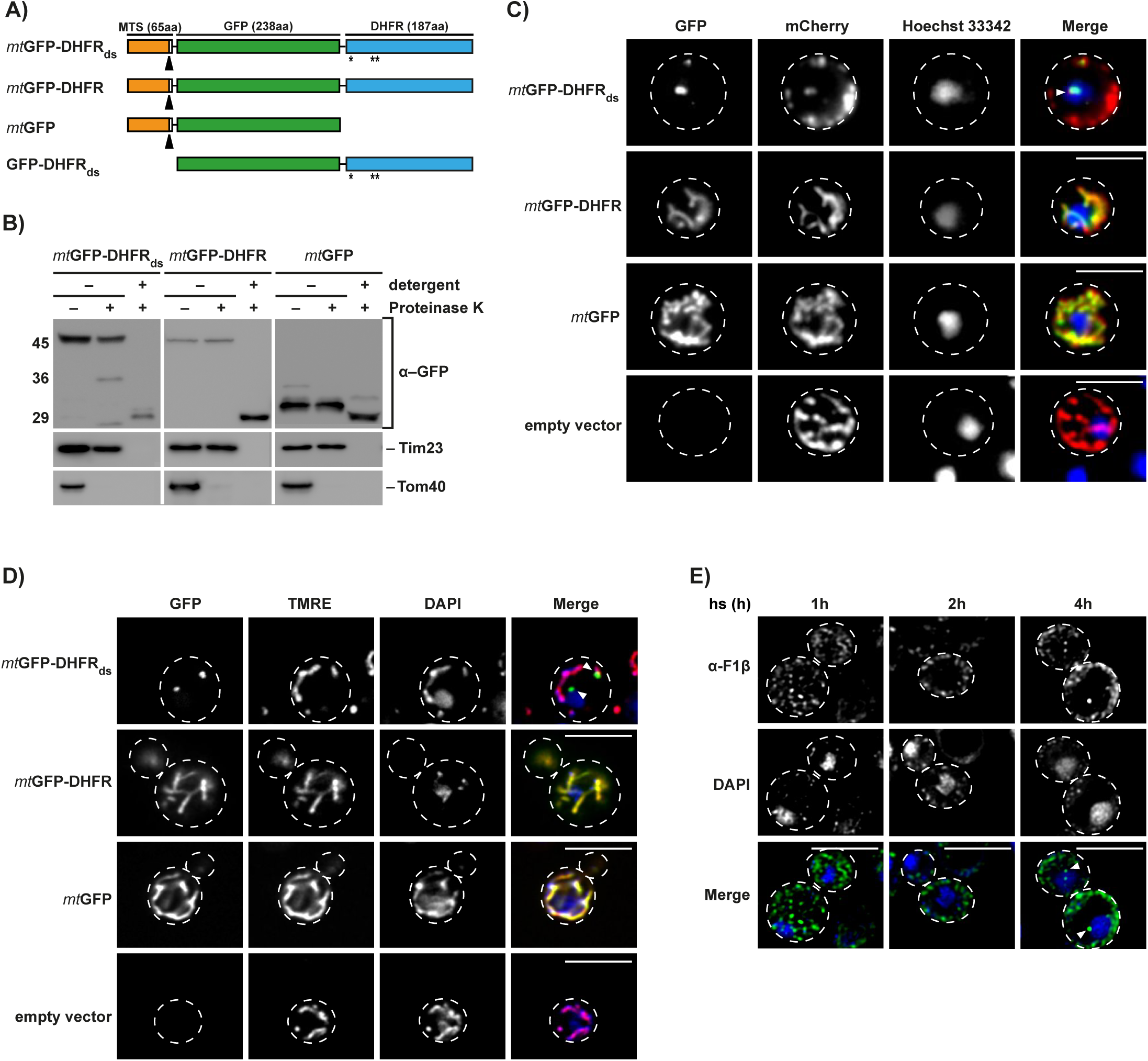
Expression and localization of aggregation-prone mitochondrial reporter proteins. A) Schematic depiction of the destabilized reporter construct *mt*GFP-DHFR_ds_ and the corresponding control constructs *mt*GFP-DHFR, *mt*GFP and GFP-DHFR_ds_. Indicated are the mitochondrial targeting sequence (orange), the GFP domain of *A. Victoria* (green) and the DHFR domain of *M. musculus*. Destabilizing mutations in the DHFR domain are indicated (*). Cleavage site for MPP is indicated by black arrow. B) Treatment of intact and lysed mitochondria containing destabilized reporter or control proteins with 2.5 µM proteinase K. Mitochondria were lysed by treatment with 0.5 % triton X-100 (detergent). Proteinase K digestion was performed for 20 min at 4 °C. Reporter proteins and control proteins of the outer (Tom40) and inner mitochondrial membrane (Tim23) were detected by Western blot. C) Analysis of aggregate containing live cells by fluorescence microscopy. Mitochondria-targeted mCherry (red) was detected in cells expressing destabilized or control proteins for 5 h or in cells containing empty vector. Nuclei were visualized by staining with Hoechst-33342 (blue). Scale bar: 10 µm. D) Cells expressing destabilized GFP-reporter or control proteins for 5 h were analyzed as above. Mitochondrial inner membrane potential (Δψ) was visualized by staining with TMRE (red) and nuclei stained with DAPI (blue). Scale bar: 10 µm. E) Immunofluorescence of yeast wild type cells upon incubation at 42°C for indicated time points. Mitochondrial protein F_1_β (green) was detected by corresponding antibodies and Nuclei were visualized by staining with DAPI (blue). Scale bar: 5 µm.

Fluorescence microscopy analysis of the control proteins *mt*GFP and *mt*GFP-DHFR revealed that they were evenly distributed within the mitochondrial network in the cell, demonstrating complete solubility. Co-localization of both *mt*GFP-DHFR and *mt*GFP with the mitochondrial marker mCherry confirmed their mitochondrial localization. In contrast, in cells expressing the destabilized *mt*GFP-DHFR_ds_ for 5 h, the reporter protein was typically concentrated into two cellular agglomerations, indicating an aggregation reaction. These agglomerations were also positive for the mitochondrial marker protein mCherry, showing that both proteins were derived from mitochondria (Fig. 1C). The microscopic analysis showed that one agglomerate was typically located within the mitochondrial network, while the second larger agglomerate contained less mitochondrial material and was separated from the mitochondrial network. We analyzed whether this separated agglomerate retained mitochondrial functionality by staining the yeast cells with TMRE, a membrane potential-dependent mitochondrial dye. TMRE staining in cells expressing stable GFP fusion proteins reflected the complete mitochondrial network, demonstrating an intact membrane potential (Fig. 1D). The separate mitochondrial agglomerate containing *mt*GFP-DHFR_ds_ aggregates showed no TMRE staining, while the remaining mitochondrial network in these cells exhibited a normal membrane potential. These findings indicate that the majority of protein aggregates formed by destabilized proteins were separated within a specialized non-functional mitochondrial compartment. To investigate, whether endogenous proteins were also sequestrated from the mitochondrial network under stress conditions, we followed the localization of the mitochondrial ATPase subunit F_1_β during incubation of yeast cells at elevated growth temperatures by immunofluorescence over time (Fig. 1E). We found that incubation at 42 °C led to fragmented mitochondria after 1 h and 2 h incubation, and resulted in the formation of bright F_1_β-containing agglomerates after 4 h stress, indicating a similar separation of damaged endogenous proteins as observed during *mt*GFP-DHFR_ds_ expression. We termed this novel mitochondrial aggregate site the <u>I</u>ntra-<u>M</u>itochondrial Protein <u>Q</u>uality Control <u>C</u>ompartment (IMiQC).

### Mitochondrial protein homeostasis is maintained in presence of destabilized proteins due to IMiQC formation

We analyzed the destabilized mitochondrial reporter protein by sedimentation assays to investigate the biochemical properties of the formed aggregates. After lysis of yeast cells under native conditions and high speed centrifugation at 20.000 × g, both *mt*GFP-DHFR and *mt*GFP were detected in the soluble fraction, while the destabilized *mt*GFP-DHFR_ds_ was found exclusively in the pellet fraction, demonstrating its complete aggregation even under normal growth conditions (Fig. 2A). Analysis of *mt*GFP-DHFR_ds_ expressing cells at different time points of expression (after 2.5 h and 10 h) showed that the destabilized protein exclusively formed large protein aggregates, pelleting already at 2.000 × g, (Fig. 2B and Sup. 2A).

**Figure 2.**
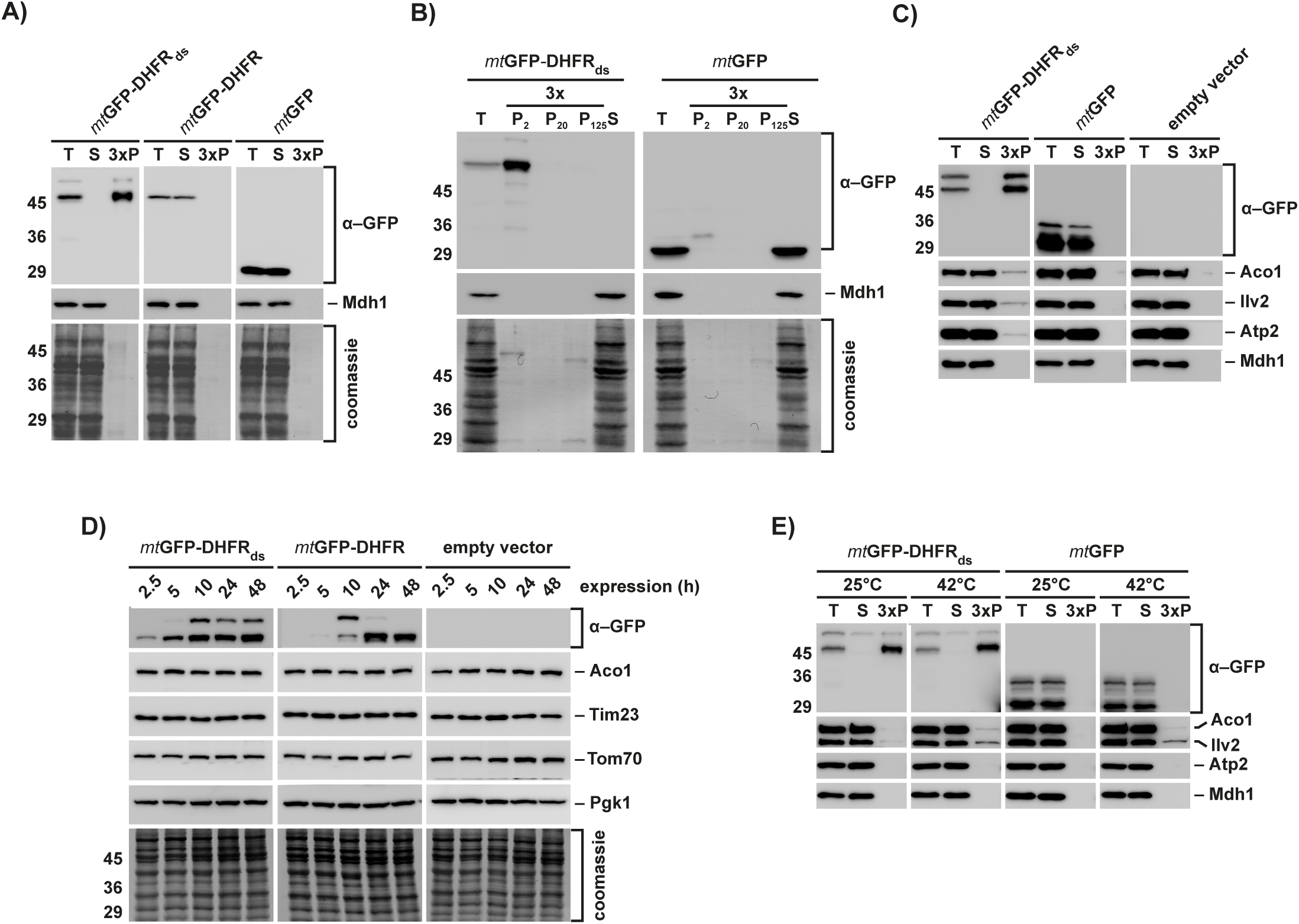
Protein homeostasis is maintained in presence of aggregated polypeptides in the mitochondrial matrix. A) The aggregation behavior of the destabilized reporter and control proteins was analyzed in wild type yeast cells after 10 h protein expression. Cells were lysed and centrifuged at 20,000 × g. Total cell extracts (T), supernatants (S) and pellet fractions (P, 3 × amounts) were applied to SDS-PAGE and analyzed by Western blot using antibodies against GFP. Mdh1 was used as soluble control protein of the mitochondrial matrix. Total protein content is indicated by the respective Coomassie-stained blot membrane sections. B) Differential centrifugation of wild type yeast cells expressing destabilized reporter protein or control proteins. Cells expressing the indicated reporter proteins for 2.5 h were lysed and pellet fractions (3 × amounts) after centrifugation at 2,000 ×g (P_2_), 20,000 ×g (P_20_), and 125,000 xg (P_125_) were analyzed by SDS-PAGE and Western blot as above. (T, total cell extract; S, supernatant after 125,000 × g centrifugation). C) Co-aggregation of endogenous proteins within the mitochondrial matrix. Cells expressing the indicated reporter proteins for 24 h were lysed and centrifuged at 20,000 × g. Supernatants (S) and pellet fractions (P, 3 × amounts) were analyzed by SDS-PAGE and Western blot as above using antibodies against the indicated proteins. D) Cells containing the indicated reporter plasmids were collected after the indicated expression times. Protein levels of the GFP-reporter proteins as well as an outer membrane protein (Tom70), an inner membrane protein (Tim23) and a mitochondrial matrix protein (Aco1) were analyzed by Western blot. The cytosolic protein Pgk1 was used as loading control. E) Mitochondria isolated from cells expressing the indicated reporter proteins for 5 h were incubated either at 25 °C or at 42 °C for 20 min. Aggregation of the reporter proteins and indicated endogenous matrix proteins was analyzed by lysis and centrifugation as described above.

To test for potential proteotoxic effects of mitochondrial protein aggregation, we analyzed a possible co-sedimentation of the mitochondrial proteins Aco1, Ilv2 and Atp2, which are already known to be thermo-unstable in yeast mitochondria ^16^. Only very minor amounts (about 1%) of the endogenous proteins co-sedimented with *mt*GFP-DHFR_ds_ aggregates after 10 h expression, increasing only slightly after long-term expression for 24 h (Fig. 2C). The major fraction of the three proteins remained soluble. As control, the thermo-stable protein malate dehydrogenase 1 was not co-aggregated with *mt*GFP-DHFR_ds_ aggregates. To test for co-aggregation in a more comprehensive manner, we analyzed the subset of mitochondrial proteins co-sedimenting with *mt*GFP-DHFR_ds_ by a 2D-PAGE approach. A general evaluation of the 2D-spot pattern revealed that all identified proteins in the aggregate fraction co-sedimented only in very minor amounts compared to the amount of the destabilized reporter protein (Sup. 3). We were able to identify 16 co-aggregating endogenous proteins. Among those identified proteins were typical metabolic enzymes, like Ilv1 and Ilv3 (involved in amino acid biosynthesis), Cit1 (TCA cycle), like, as well as proteins of the mitochondrial inner membrane, which are part of the electron transport chain (Atp1 and Atp2). To assess the cellular reactivity to the expression of destabilized mitochondrial proteins, we followed the protein levels of mitochondrial proteins of the matrix, IMM as well as OMM over 48 h upon expression of the reporter proteins (Fig. 2D). The amounts of all tested proteins were stable during the experimental time frame, excluding changes in mitochondrial protein biogenesis or turnover due to proteotoxic stress.

We tested the sedimentation behavior of the endogenous mitochondrial proteins also under heat stress conditions. We incubated isolated mitochondria at 42 °C for 40 min and analyzed a co-aggregation with *mt*GFP-DHFR_ds_. Although Aco1 and Ilv2 started to aggregate at higher temperatures as expected, the amount of aggregated proteins was similar in mitochondria containing the misfolded *mt*GFP-DHFR_ds_ (Fig. 2E). This result suggested that even under additional stress conditions, mitochondrial aggregation processes were not excessively increased. These observations indicated that the accumulation of destabilized proteins IMiQC, is able to neutralize misfolded protein species and contributes significantly to protein homeostasis of mitochondria.

### Mitochondrial protein quality control system is relieved by sequestration of destabilized proteins into IMiQC

To analyze if components of the mitochondrial PQC system were recruited to IMiQC, we assayed aggregate contents after long-term expression of *mt*GFP-DHFR_ds_ and control proteins. Hsp78, the mitochondrial ClpB-type chaperone as well as the matrix AAA+ protease Pim1 did not co-sediment with the protein aggregates In contrast, as we already observed by 2D-PAGE analysis, a small amount of the mitochondrial Hsp70-type chaperone Ssc1 could be found in the pellet fraction, its amounts slightly increasing from 5 h to 24 h expression. The mitochondrial Hsp60 chaperone was found in the pellet fraction after 10 h and 24 h (Fig. 3A; Sup. 3). These findings indicate, that Ssc1 as well as Hsp60 interact with mitochondrial aggregates. However, we estimated that only about 1% of the total protein amount of both Ssc1 and Hsp60 present within mitochondria was found associated with the destabilized reporter proteins.

**Figure 3.**
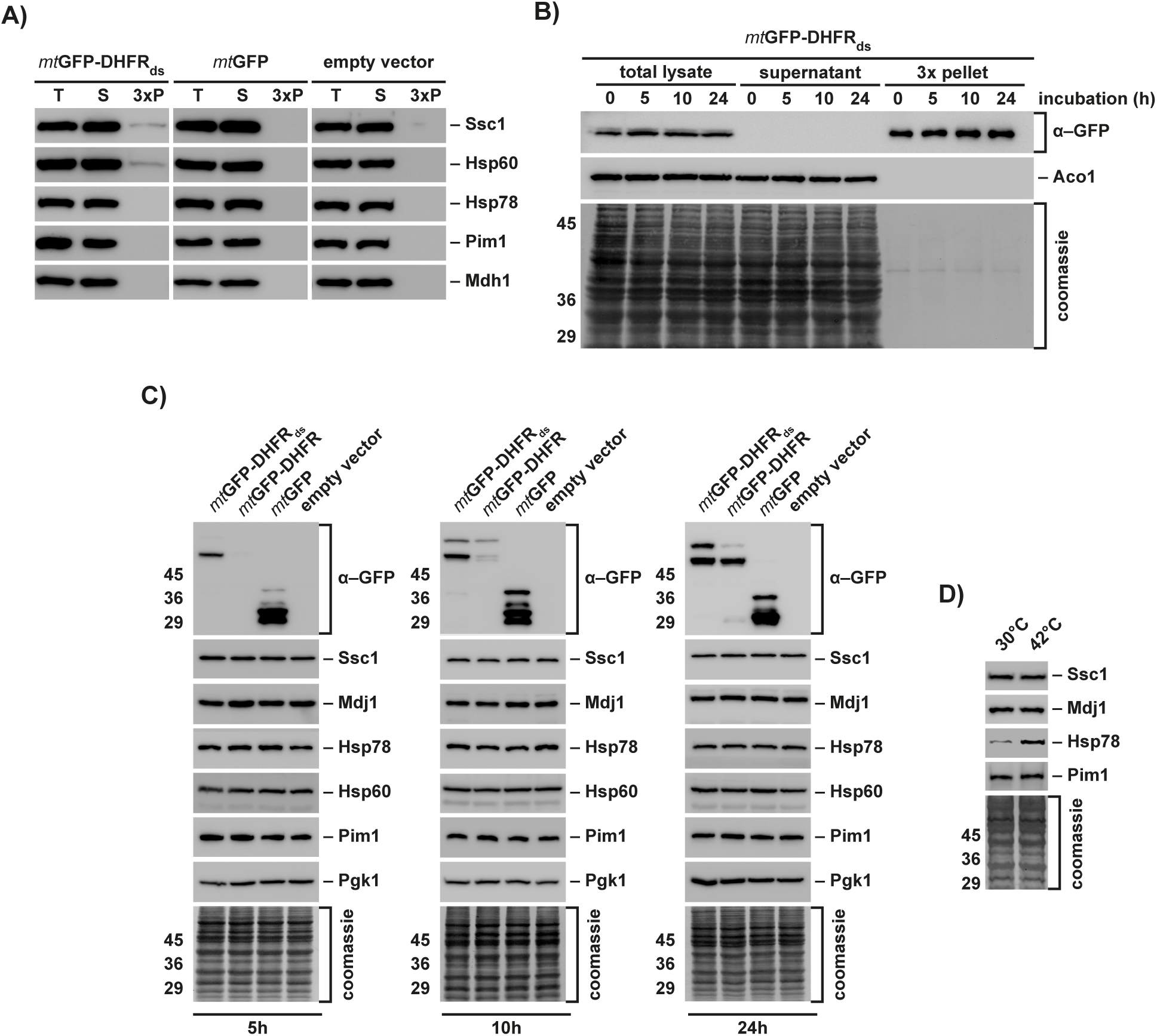
Components of the mitochondrial PQC system are not affected by the accumulation of aggregated proteins. A) Co-aggregation of mitochondrial PQC system components. Wild type yeast cells expressing reporter proteins for 24 h were lysed and centrifuged at 20,000 × g. Total cell extracts (T), supernatants (S) and pellet fractions (P, 3 × amounts) were analyzed by Western blot using antibodies against GFP and the indicated mitochondrial PQC components. B) Long-term stability of IMiQC-Aggregates. Expression of *mt*GFP-DHFR_ds_ was blocked after 2.5 h by addition of 2 % glucose. Cells were further incubated at 30 °C for the indicated times. Mitochondrial aggregates were analyzed by Western blot as described. Aco1 was used as control protein. C) Protein levels of mitochondrial PQC components in cells expressing destabilized reporter or control proteins for 5 h, 10 h or 24 h. Total cell extracts were analyzed by Western blot using the indicated antibodies. The cytosolic protein Pgk1 was used as loading control. D) Protein levels of mitochondrial PQC components in cells incubated at 30 or 42 °C for 4 h were analyzed by Western blot as above.

We also tested the long-term stability of the mitochondrial aggregates. We stopped the expression of *mt*GFP-DHFR_ds_ by addition of glucose after 2.5 h and incubated the cells additionally for up to 24 h (Fig. 3B). Even 24 h, *mt*GFP-DHFR_ds_ remained in the aggregate pellet. The protein amounts were very similar to the beginning of the experiment, indicating that the IMiQC inclusion was neither resolubilized nor degraded in the tested time frame

The accumulation of misfolded polypeptides under the used conditions would be also able to elicit an UPR^mt^ reaction. We determined the protein levels of several mitochondrial PQC components after different time points of *mt*GFP-DHFR_ds_ expression and found that even after 24 h none of the mitochondrial members of the PQC system were upregulated, demonstrating that the presence of mitochondrial aggregates was not able to trigger a heat stress response equivalent (Fig. 3C). As a control, we performed a heat stress experiment by incubating cells for 4 h at 42 °C. As expected, under heat stress, ClpB-type chaperone Hsp78 showed a significant upregulation.

We conclude that IMiQC formation effectively sequestered the potentially hazardous protein species, thereby preventing occupation and subsequent overload of the mtPQC system.

### Mitochondrial health and functionality are maintained by IMiQC formation

We performed a comprehensive analysis of important mitochondrial processes to assess the impact of expressed *mt*GFP-DHFR_ds_. To investigate growth phenotype, cells were grown either on fermentable glucose medium or on non-fermantable glycerol medium. In both cases, cells expressing *mt*GFP-DHFR_ds_ grew like control cells, indicating that destabilized proteins within mitochondria had neither toxic effects on cellular nor on mitochondrial processes (Fig. 4A). We measured the membrane potential of these mitochondria using the fluorescent dye DiSC_3_, which is quenched in a membrane potential dependent manner ^17^. Recovery of DiSC_3_-fluorescence upon addition of sodium azide was similar in mitochondria containing *mt*GFP-DHFR_ds_ compared to control mitochondria (Fig. 4B), indicating that the mitochondrial membrane integrity and respiratory efficiency were not affected by the presence of destabilized proteins. We also examined the ability of mitochondria to import precursor proteins from the cytosol. Protein import requires a coordinated function of many mitochondrial systems, like import machinery, respiratory chain and chaperones. The *in vitro* translated radiolabeled preprotein Su9(70)-DHFR was imported *in vitro* into mitochondria isolated from yeast strains expressing the destabilized *mt*GFP-DHFR_ds_ or control proteins. Su9(70)-DHFR was efficiently imported into mitochondria irrespective of the presence of aggregated proteins, indicating a full activity of the preprotein import machinery (Fig. 4B). To determine whether mitochondrial aggregation interfered with proteolytic processes, we performed degradation assays of an *in vitro* imported radiolabeled reporter protein b_2_(167)Δ-DHFR. The degradation efficiency in mitochondria isolated from aggregate-containing cells was similar to control mitochondria, indicating that due to IMiQC formation the overall PQC-relevant degradation processes in the mitochondrial matrix remained intact (Fig. 5A). A structural analysis of mitochondrial protein complexes of the oxidative phosphorylation system, TCA cycle as well as the preprotein translocase of the outer membrane (TOM) by BN-PAGE did not show any alterations caused by a presence of protein aggregates (Sup. 3). Taken together, the experiments indicate that mitochondrial protein homeostasis was not affected by the accumulation of misfolded proteins.

**Figure 4.**
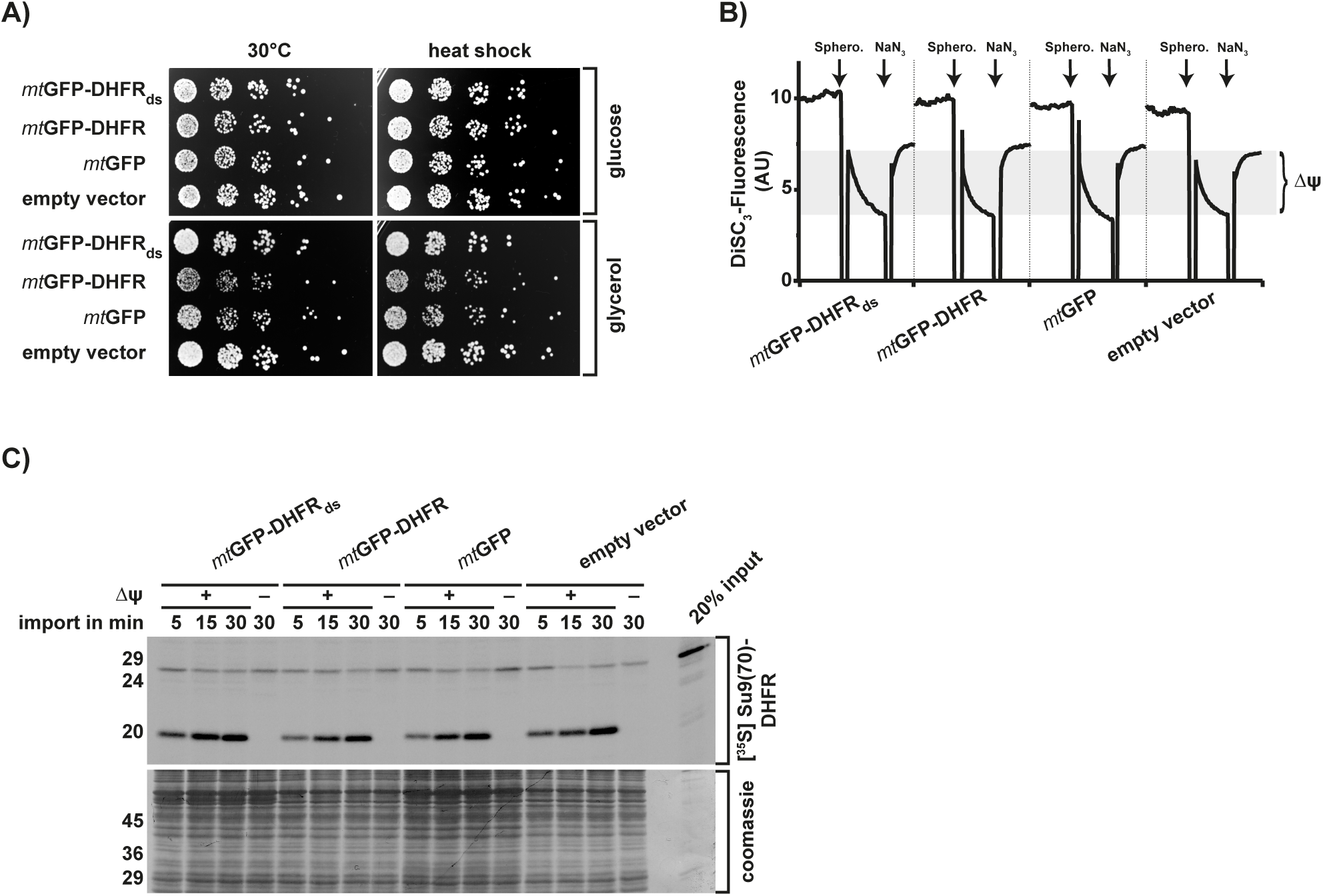
Mitochondrial quality and functionality are maintained in cells containing IMiQC-aggregates. A) Growth of *mt*GFP-DHFR_ds_ expressing cells was analyzed. After 10 h expression of indicated reporter proteins, cells were incubated at 30 or 42 °C (heat shock) for 1 h and grown on fermentable (2 % glucose) or non-fermentable (3 % glycerol) medium at 30 °C. B) Determination of mitochondrial membrane potential (Δψ) in yeast wild type cells. Spheroblasts were prepared from cells expressing destabilized reporter or control proteins for 10 h and analyzed using DiSC_3_ fluorescence as described in Experimental procedures. C) *In vitro* import of cytosolic precursor proteins. Radiolabeled Su9(70)-DHFR was imported for indicated time points into isolated and energized mitochondria containing *mt*GFP-DHFR_ds_ or control proteins (10 h expression) as described in experimental procedures. Where indicated (−Δψ), the mitochondrial membrane potential was abolished by addition of 8 µM antimycin A, 0.5 µM valinomycin and 2 µM oligomycin. Import reactions were analyzed by SDS-PAGE and digital autoradiography.

### Mitochondrial aggegation influences functional aspects of the mitochondrial protease Pim1

Previous experiments in our group showed a prominent involvement of the matrix AAA+ protease Pim1 in the removal of polypeptides damaged by reactive oxygen species (ROS) ^18^. When we performed cellular growth analysis in presence of the ROS-inducing chemical menadione, cells expressing *mt*GFP-DHFR_ds_ showed an increased ROS resistance after 5 h expression of the destabilized polypeptide. However, cells became increasingly sensitive to ROS after longer expression times, resulting in a significantly decreased ROS resistance after 24 h expression (Fig. 5B). Notably, cells expressing the control proteins *mt*GFP-DHFR and *mt*GFP showed the opposite pattern. We confirmed these observations by performing the ROS resistance analysis at an elevated culture temperature of 37 °C and found a similar behavior for cells containing *mt*GFP-DFHR_ds_ aggregates. The increased ROS-sensitivity of *mt*GFP-DHFR_ds_ expressing cells indicated that mitochondrial aggregate formation influenced Pim1-related functions. To directly test the involvement of Pim1, we performed the ROS resistance assay in Pim1 overexpressing cells in presence or absence of aggregates. Indeed, elevated levels of the protease Pim1 completely rescued the increased ROS sensitivity phenotype of cells expressing *mt*GFP-DHFR_ds_ even at elevated temperatures.

**Figure 5.**
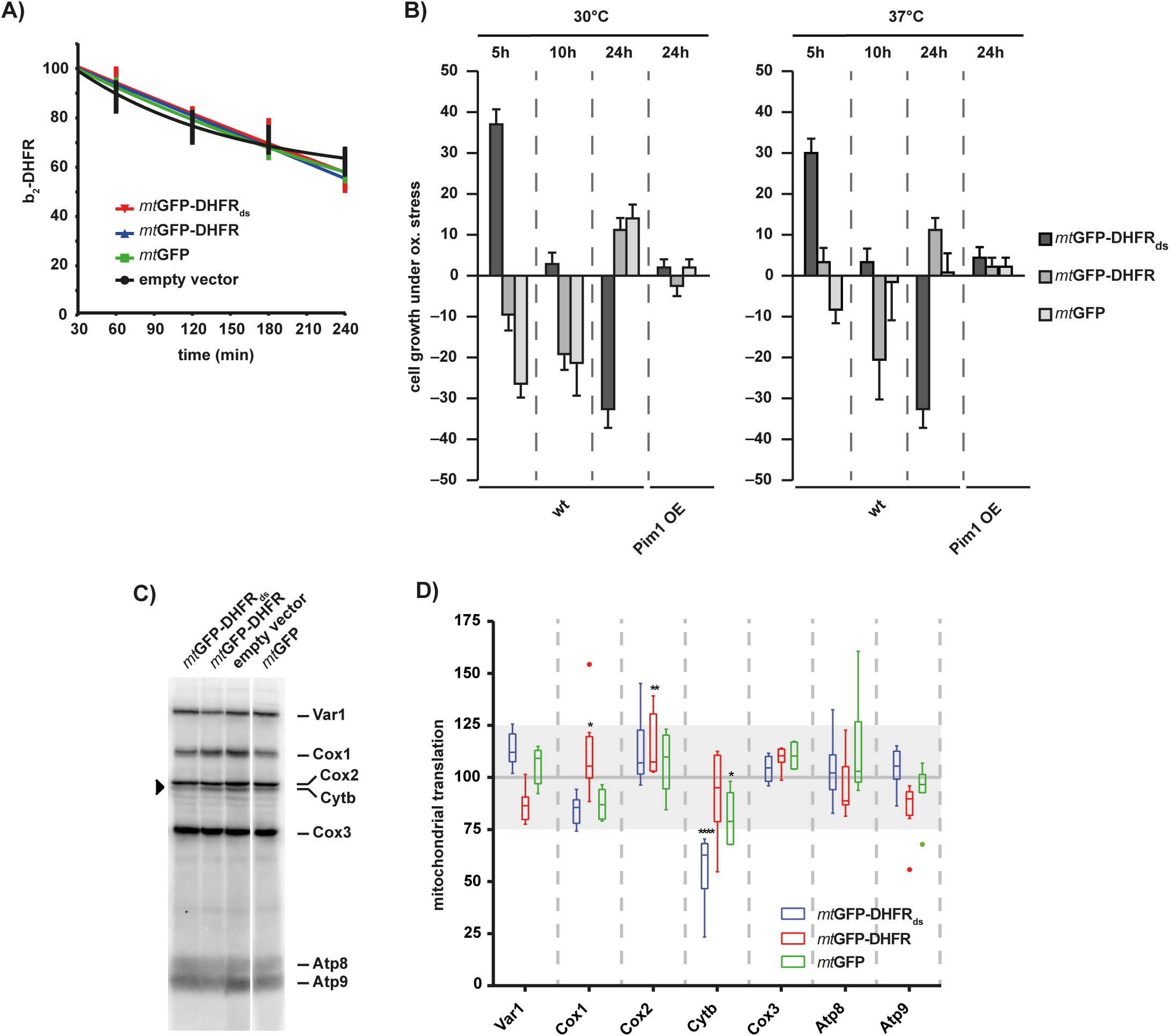
IMiQC interferes with Pim1 function. A) Protein degradation in the mitochondrial matrix. Radiolabeled *cytb_2_*(167)Δ-DHFR was imported into the matrix of isolated mitochondria expressing the indicated reporter proteins for 10 h for 20 min. After the import reaction, samples were withdrawn at the indicated time points and analyzed by SDS-PAGE and autoradiography. Residual protein levels were quantified using MultiGauge (Fujifilm). Mean values and SDm bars (n=5) are shown..B) ROS resistance of IMiQC containing cells. Wild type or Pim1-overexpression (OE) cells expressing destabilized reporter or control proteins for the indicated times were grown on fermentable minimal medium (2 % glucose) in presence of 20 mM menadione at 30 or 37 °C. Growth inhibition zones were measured and quantified as resistance to menadione relative to control cells. 0% represents cells containing empty vector, 100% represents complete resistance to menadione.. Shown are mean values and SDm (n=5). C) and D) *In organello* translation rates. Isolated mitochondria containing destabilized reporter or control proteins (10 h expression; indicated in colors) were incubated with [^35^S]-methionine/cysteine for 45 min at 30 °C (4.4 µCi/ reaction). Translation products were analyzed by 15 % Urea-SDS-PAGE and detected by digital autoradiography. D) Quantification of three independent experiments (representative autoradiogram shown in C). The values were normalized to protein levels of translation products in mitochondria isolated from cells containing empty vector. Quantifications are shown as a box-whisker diagram. Grey area indicates fluctuation range between individual translation experiments.

Pim1 was previously described to participate also in mitochondrial gene expression, independently of its proteolytic activity ^19^. To analyze whether mitochondrial gene expression was affected by the presence of mitochondrial aggregates, we performed an *in organello* translation assay, in which newly synthesized proteins were labeled by the incorporation of [^35^S]-methionine/cysteine in isolated mitochondria. After separation of the translation products we detected 7 of 8 mitochondria-encoded proteins (Fig. 5C and D). Although the *in organello* translation efficiencies of the individual polypeptides fluctuated for about 25% in repeated experiments, the translation of cytochrome *b* was significantly reduced in *mt*GFP-DHFR_ds_ containing mitochondria. None of the other mitochondria-encoded proteins did show significant alterations in translation efficiency. As Pim1 was shown to be specifically required for the maturation of cytochrome *b* mRNA in yeast mitochondria ^19^, this observation confirms a negative influence of IMiQC accumulation on Pim1 function.

### IMiQC is stored at the nucleus to allow asymmetric distribution during budding

We analyzed the number of mitochondrial aggregates during *mt*GFP-DHFR_ds_ expression over time. Yeast cells were either incubated in presence of galactose to obtain a continuous expression or alternatively, glucose was added after 2.5 h to repress further expression. After a subsequent incubation for 5-24 h, the number of GFP-positive dot structures per cell were counted (Fig. 6A and 6B). After 5 h expression, each cell contained on average 3 aggregate deposits. During constitutive expression of *mt*GFP-DHFR_ds_, the number of aggregate structures increased over time, leading to numbers between 4 and 6 dots per cell after 24 h. Upon inhibition of expression, the number of aggregates decreased over time, resulting in only one or two dots per cell after 24 h, while total protein levels of the expressed constructs remained similar over 24 h (Fig. 3B). These results suggest that misfolded proteins form first several smaller aggregate structures within mitochondria, which fuse at longer incubation times to one or two large aggregate compartments.

**Figure 6.**
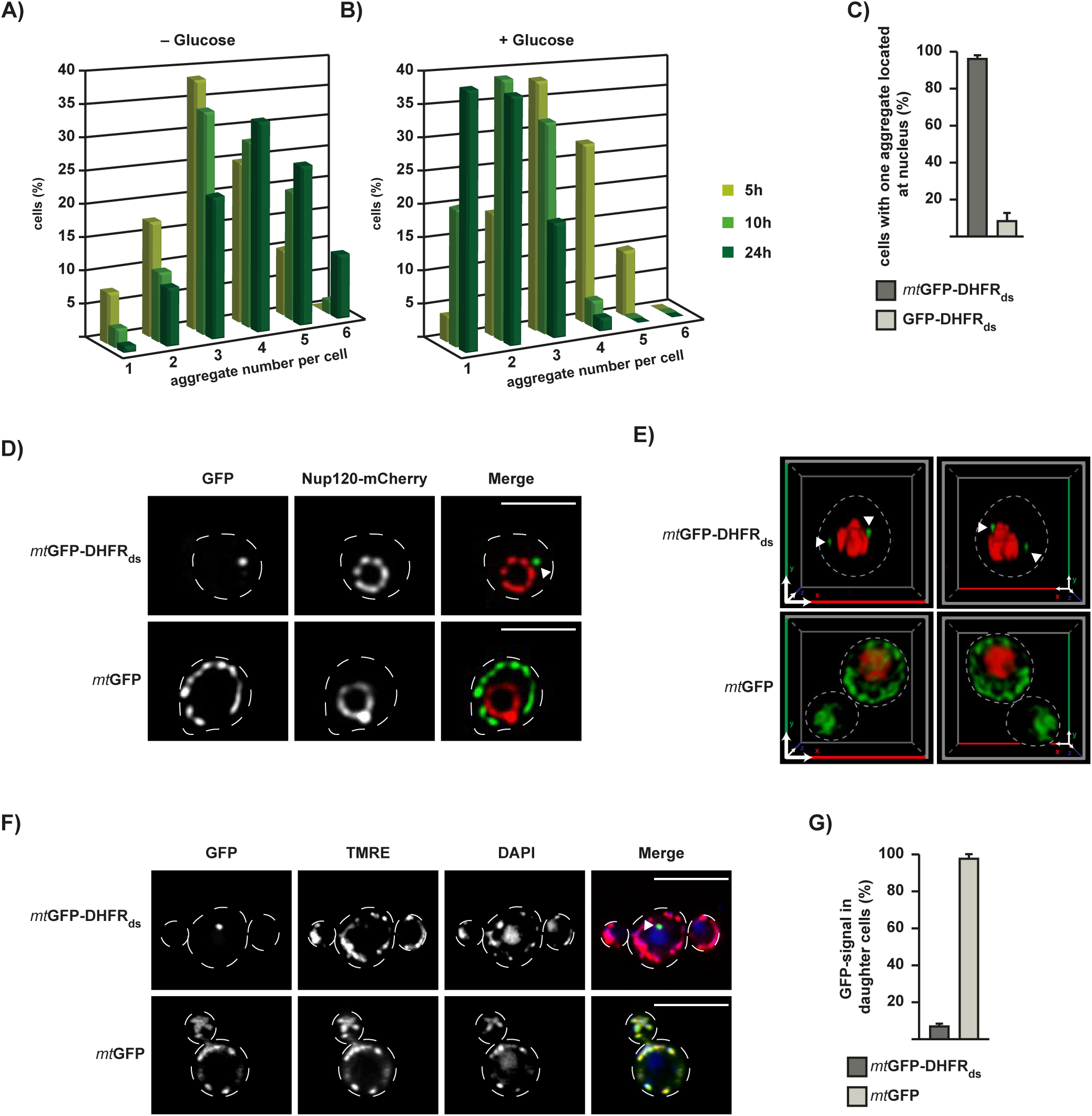
Formation of IMiQC leads to aggregate sequestration at the nucleus. A) Quantification of aggregate number in cells expressing *mt*GFP-DHFR_ds_. Expression was either continuous up to 24 h (left panel) or stopped after 2.5 h by addition of 2 % glucose (right panel) or remained constant. Live cells were analyzed by fluorescence microscopy and GFP-positive dots were counted in 200 cells each after 5 h (light green), 10 h (green) and 24 h (dark green). The entirety of all expressing cells was set to 100%. B) Quantification of aggregate localization. Cells expressing *mt*GFP-DHFR_ds_ (dark grey) or cytosolic GFP-DHFR_ds_ (light grey) for 5 h that contained mitochondrial aggregates located in the nuclear periphery were counted (100 cells each, set to 100 %). Mean values and SDm bars are shown (n=3). C) Peri-nuclear localization of IMiQC. Yeast cells expressing the nuclear pore protein Nup120-mCherry (red), *mt*GFP-DHFR_ds_ or the control protein *mt*GFP (green) for 5 h were analyzed by fluorescence microscopy. Scale bar: 5 µm. D) 3-dimensional reconstruction of IMiQC-aggregates (green, upper panel) or mitochondria-located GFP (green, lower panel) in yeast cells containing Nup120-mCherry (red). 3-D reconstruction was simulated after Z-stack scan using Fiji. F) Distribution of IMiQC structures during cell division. Fluorescence microscopy of live yeast cells expressing *mt*GFP-DHFR_ds_ or the control protein *mt*GFP (green) for 5 h. Expression of reporter proteins was stopped by addition of 2 % glucose and cells were incubated at 30 °C for additional 3 h. Mitochondria were visualized by TMRE-staining (red) and nuclei by DAPI-staining (blue). Scale bar: 10 µm. G) Quantification of yeast cell buds containing aggregate structures. Cells expressing indicated reporter proteins were analyzed by fluorescence microscopy as above. 100 daughter cells during budding process were analyzed for the presence of GFP-positive dots in three independent experiments. Mean and SDm bars are shown.

To investigate whether the separated aggregate was residing inside or localized just close to the nucleus, we co-expressed *mt*GFP-DHFR_ds_ with mCherry-tagged Nup120, a protein of the nuclear pore complex. While, as already shown, *mt*GFP localization reflected the mitochondrial network and was not located at the nucleus, the peri-nuclear aggregates formed by *mt*GFP-DHFR_ds_ were clearly located close to but not inside the nucleus (Fig. 6C). This observation was confirmed by a 3-dimensional reconstruction of the cellular IMiQC localization (Fig. 6D). About 96% of all cells expressing *mt*GFP-DHFR_ds_ showed at least one aggregate located at the nucleus. We expressed a destabilized GFP-DHFR_ds_ fusion protein, which did not contain a MTS, in the cytosol. Although these cells also showed prominent cytosolic aggregate structures visible as single dots, a peri-nuclear localization was only observed in about 8% of all cells analyzed (Fig. 6E), These observations indicate that a peri-nuclear localization of mitochondrial aggregate structures is a specific feature of the IMiQC.

To test if the IMiQC is distributed to daughter cells during cell division, we analyzed cells expressing either *mt*GFP or *mt*GFP-DHFR_ds_ for 5 h, stopped the expression by addition of glucose and then incubated the cells for additional 3 h. Non-aggregating *mt*GFP was distributed to the daughter cells in 98% of all expressing cells, correlating with a normal distribution of mitochondria from the mother cell to the bud (Fig. 6F and G). In contrast, while TMRE staining of mitochondria revealed a normal propagation of mitochondria per se, only 7% of cells expressing *mt*GFP-DHFR_ds_ exhibited aggregate structures in the daughter cells. This observation strongly argues that the IMiQC was not distributed to daughter cells, resulting in an eventual elimination of polypeptide aggregates during multiple rounds of cell division.

### Detoxification of aggregates and IMiQC formation requires fission and fusion of mitochondria

To characterize which cellular factors are involved in IMiQC formation, we analyzed different mutant yeast strains lacking mitochondrial chaperones. We also tested the effect of factors required for the regulation of mitochondrial dynamics. First, we compared the growth rates of the respective mutant strains expressing the destabilized reporter *mt*GFP-DHFR_ds_ or its control proteins. On fermentable and non-fermentable carbon sources, we found that cellular growth rates were not affected by aggregate formation (Fig. 7A). Even in mutant cells, where members of the mitochondrial Hsp70 system were deactivated (*ssc1-3* and *mdj1*∆), as well as in *hsp78*∆ cells, no significant growth differences were observed. This supports our conclusion that IMiQC formation allows a functional detoxification of damaged proteins independently of the mitochondrial PQC system. Off note, *mdj1* deletions strains are respiratory deficient, preventing the growth analysis on non-fermentable carbon sources.

**Figure 7.**
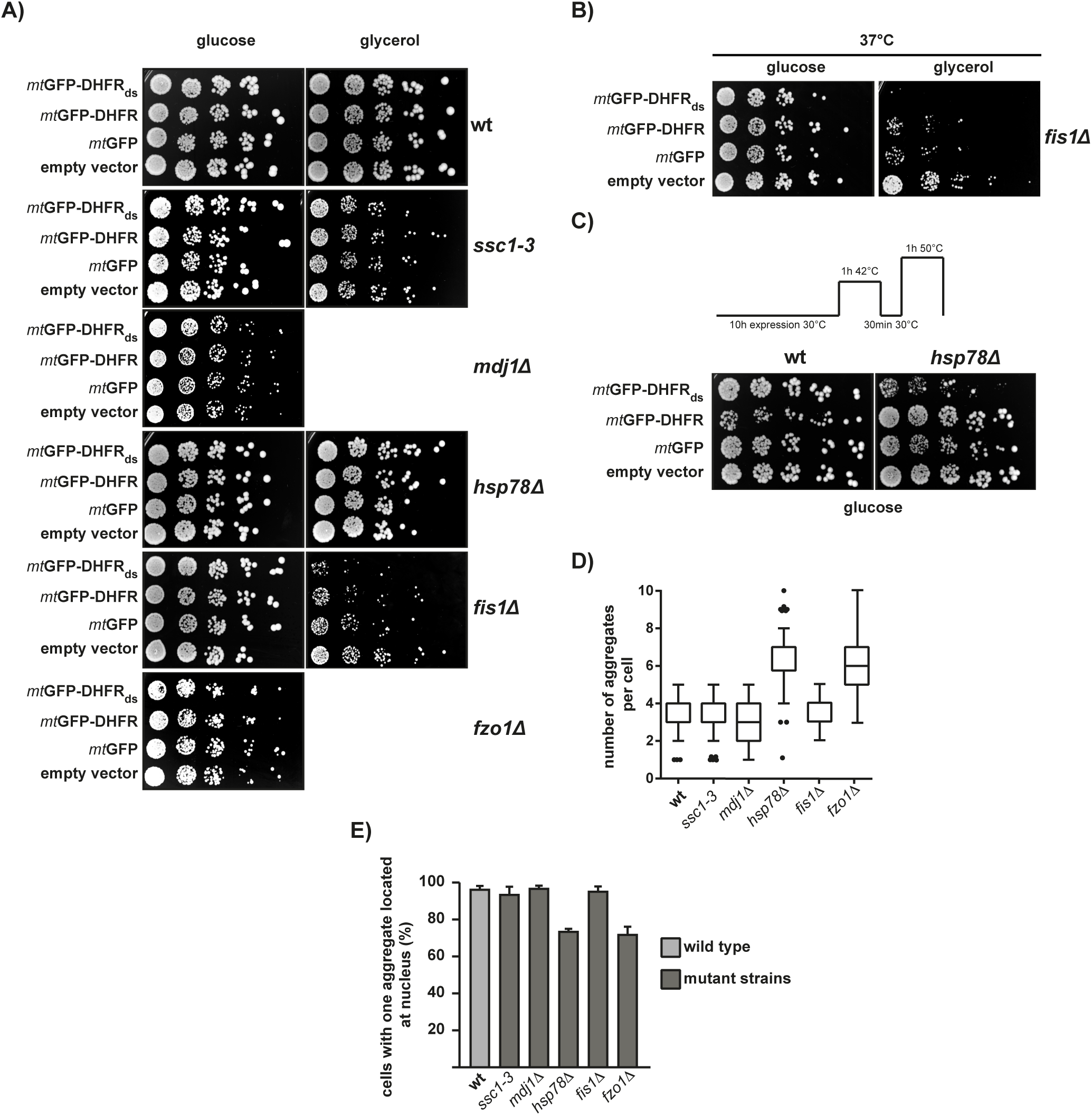
Cellular factors involved in IMiQC formation. A) Growth rates of mutant yeast cells expressing *mt*GFP-DHFR_ds_ or control proteins for 10 h. Cells from the indicated mutant strains were spotted on selective plates containing fermentable (2 % glucose) or non-fermentabe carbon source (3 % glycerol) and incubated at 30 °C. The deletion strains *mdj1∆* and *fzo1∆* are respiration deficient and do not grow on non-fermentable carbon sources. B) Cellular growth of *fis1-*cells expressing the destabilized reporter protein for 10 h under mild thermal stress. Cells were spotted as described above and incubated at 37 °C. C) Analysis of acquired thermotolerance in wild type and *hsp78∆* deletion cells expressing *mt*GFP-DHFR_ds_ or control proteins for 10 h. After protein expression at 30 °C, cells were subjected to a short heat shock and recovery as indicated. Cells were then incubated at a lethal temperature of 50 °C for 1 h and spotted on selective plates containing 2 % glucose. D) Aggregate number in yeast mutant cells. The indicated mutant cells expressing *mt*GFP-DHFR_ds_ were analyzed by fluorescence microscopy. GFP-positive aggregate dot numbers were evaluated in 200 cells each and shown as box-whisker diagram. E) Localization of aggregates in yeast mutant cells. A peri-nuclear localization of aggregate structures was analyzed in the indicated mutant strains as described in Fig. 6B. For each strain, 100 cells were counted and set to 100 %. Shown are mean values and SDm (n= 3).

Growth analysis of a knockout strain lacking the mitochondrial fission factor Fis1 led to significant a growth defect of cells containing aggregates on a non-fermentable carbon source, while growth on fermentable carbon source was similar to control cells (Fig. 7A). When we increased the growth temperature of *fis1*∆ cells to 37 °C, the expression of the destabilized polypeptides caused a lethal phenotype on non-fermentable medium (Fig. 7B). Deletion cells of the fusion factor Fzo1 expressing *mt*GFP-DHFR_ds_ grew like control cells and showed no aggregate toxicity of the mitochondrial aggregates. Unfortunately, the *fzo1*∆ cells are also respiratory deficient and could not the analyzed under non-fermentable conditions. These results indicate that the absence of a functional fission process leads to a severe perturbation of mitochondrial activity and to a complete abrogation of the protective effect of IMiQC formation, in particular under cellular stress conditions. We also tested the thermotolerance of *hsp78*∆ cells in presence and absence of aggregate accumulation (Fig. 7C). In wild-type cells, the phenomenon of acquired thermotolerance allows the survival at the lethal temperature and was not influenced by the presence of misfolded proteins in mitochondria. However, *hsp78*∆ cells expressing the destabilized *mt*GFP-DHFR_ds_ became strongly sensitive to heat stress under these conditions, indicating a role of Hsp78 in detoxification of unfolded proteins via IMiQC formation.

We also analyzed the number of aggregate structures per cell in the mutant strains. While the number of aggregates was not altered in cells lacking Fis1 compared to wt cells, it was significantly increased in *fzo1*∆ cells (Fig. 7D). Notably, the aggregate number was also increased in *hsp78*∆ cells. The microscopic analysis revealed that also the peri-nuclear localization of IMiQCs was affected in *fzo1Δ* and *hsp78Δ* cells (Fig. 7E). We conclude that a fusion of aggregate-containing mitochondrial particles is necessary for the formation of the typical peri-nuclear IMiQC. In addition, the chaperone Hsp78, although not co-sedimenting with the aggregated polypeptides, seems to be required for the formation of the large aggregated structures and influences the protective effect of the IMiQC.

## Discussion

Sequestration of protein aggregates into distinct subcellular compartments has been recently described as a specific protein quality control pathway that reduces the proteotoxic potential of an accumulation of destabilized or misfolded polypeptides ^5,6^. Here we describe a novel quality control compartment for destabilized mitochondrial proteins in *Saccharomyces cerevisiae* and the first for endosymbiontic organelles. We termed this compartment IMiQC (for <u>I</u>ntra-<u>M</u>itochondrial Protein <u>Q</u>uality <u>C</u>ompartment) where aggregation-prone mitochondrial proteins are accumulated and sequestered in a specific, mitochondria-derived compartment localized at a distinct site inside the cell. IMiQC formation essentially maintains mitochondrial homeostasis and prevents the proteotoxic impact of destabilized polypeptides on mitochondrial functions. Thus in addition to INQ, CytoQ and IPOD, IMiQC represents a fourth protein quality control compartment identified in yeast. Deposition of aggregates in distinct cellular locations (sequestration) is a secondary defense response to proteotoxic stress and takes place when the protein quality control system as a primary defense level, consisting of molecular chaperones and proteases, is not functional or overwhelmed. Notably, we observed IMiQC formation not only in presence of the destabilized reporter proteins, but also under severe stress conditions with endogenous mitochondrial proteins like the mitochondrial ATPase subunit F_1_β. To our knowledge, other aggregate compartments described so far were either observed only after overexpression of reporter proteins or under inhibition of PQC components.

As indicated by our phenotypic analysis of cells expressing the destabilized reporter proteins, IMiQC formation relieves the mitochondrial PQC system from potential proteotoxic polypeptides to increase mitochondrial fitness. In our system, IMiQC formation was observed in presence of a fully functional mitochondrial PQC system, indicated by normal mitochondrial protein degradation rates. A similar cyto-protective role of aggregate sequestration has been already observed in the case of the of cytosolic Q-body formation ^9^. Mutants of the chaperone system, e.g. *hsp104*∆ as well as *hsp42*∆ exhibited a restricted formation of aggregate deposits showed a declined viability when exposed to heat stress. In case of mitochondrial protein aggregation, IMiQC formation also protected the metabolic activity of the majority of mitochondria in the affected cells from toxic effects of destabilized proteins. The membrane potential of the part of the mitochondrial network not containing aggregates remained largely intact while the membranes surrounding the IMiQC did not exhibit a potential. As an analysis of cell growth at different temperatures showed, aggregate accumulation did not negatively affect mitochondrial fitness even in presence of additional stress. Taken together, mitochondria remained surprisingly resilient to the effects of aggregate accumulation. We conclude from our observations that aggregate sequestration in form of IMiQC is a major factor in the resistance to proteotoxic impacts.

Correlating with the absence of major functional defects, we observed only a very minor extent of co-aggregation between the destabilized reporter proteins and other endogenous proteins components in the mitochondria. Even under heat stress conditions, no increased co-aggregation of relatively aggregation-prone proteins like Aco1 and Ilv2 ^16^ was observed. In general, the amounts of these polypeptides in the soluble fractions did not change in a detectable manner, indicating a largely sustained enzymatic activity. A similar behavior was observed for typical members of the mitochondrial PQC machinery. In other systems a significant amount of molecular chaperones, in particular of the Hsp70 and Hsp100 families, was found associated with aggregate deposits ^3, 5^. In contrast, we observed only a minor titration of matrix chaperones to the destabilized reporter proteins, indicating that the protein folding capacity remains essentially normal. Indeed, import of cytosolic preproteins into the matrix, a reaction that is absolutely dependent on the activity of the mitochondrial Hsp70 system was not affected at all. As there was neither an obvious change in the soluble amounts of Hsp60 and mtHsp70, nor any indication of a mitochondrial UPR reaction we conclude that IMiQC formation prevented the triggering of a mitochondrial stress response. The absence of a disaggregation reaction also indicated that the IMiQC is a stable and essentially inert compartment. Previous experiments from our group have indicated that mtHsp70 is mainly involved in prevention of aggregation at physiological temperatures, but not at elevated temperatures ^16^. Hence, it is conceivable that under long-term heat stress conditions other protective reactions, like IMiQC formation, becomes more relevant. Indeed, our results about the sequestration of the endogenous mitochondrial matrix protein F_1_β revealed that IMiQC formation took place only after several hours of stress, while mitochondrial fragmentation was observed already at early time points. Moreover, *hsp78*∆ cells exhibited a strong defect of acquired thermotolerance when the destabilized reporter protein was expressed. Hsp78 function seems to be necessary for IMiQC formation since the respective deletion cells contained a significantly larger number of smaller aggregates that were not able to merge to the typical IMiQC. A similar effect was already observed during the formation of cytosolic protein deposits, in which deletion of Hsp104 led to a block in the progression of aggregate deposition, resulting in an accumulation of multiple peripheral protein aggregates ^9^.

In yeast mitochondria the ATP-dependent protease Pim1 is the only matrix protease that preferentially recognizes and degrades unstructured polypeptides ^20, 21^. Pim1-mediated degradation of damaged proteins becomes particularly important under oxidative stress conditions that do not allow refolding to the active conformation ^18, 22^. Interestingly, Pim1 seemed to be the only mitochondrial PQC component that was specifically affected after expression of aggregation-prone polypeptides. While its overall proteolytic activity did not seem to be altered by the presence of aggregated polypeptides, a careful phenotypic analysis demonstrated a strong and specific defect in mitochondrial translation of cytochrome *b* and an increased sensitivity of cells to long-term treatment with ROS-generating chemicals. Both observations indicate a functional defect of Pim1 as biogenesis of cytochrome *b* requires its activity ^19^ and *pim1*∆ cells were shown to be more sensitive to ROS treatments ^18^. Taken together we conclude that the reduction of the load of misolded proteins on the PQC system by IMiQC formation results in only subtle defects in Pim1 functionality.

Interestingly, the IMiQC represented a very stable compartment. Even after longer incubation periods, we did neither observe a resolubilization of the IMiQC-sequestered polypeptides nor a degradation by the mitochondrial PQC system even after extended post-incubation times. However, as far as data are available, polypeptides accumulated in amorphous aggregates like CytoQ and INQ can be resolubilized and degraded by the cellular PQC machinery ^5^. In case of aggregated polyQ-containing proteins, accumulated as IPOD, a removal by autophagy has been shown in yeast cells, requiring the proteins Cue5 and Atg8 ^5, 23^. Similarly, in mammalian cells a degradation of proteins collected in aggresomes by autophagy has been demonstrated ^24, 25^. Our observation of unchanged aggregate amounts at least up to 48 h incubation would argue against a clearance process similar to stress-induced mitophagy as has been observed in mammalian cells ^14^.

A major advantage of the localization of aggregated polypeptides to distinct deposition sites is the possibility to get rid of the aggregates by an asymmetric distribution during cell division. All the different cellular aggregate compartments analyzed so far showed this behavior ^26–29^. Indeed, the IMiQC aggregates were also retained in the mother cells during the budding process, essentially producing aggregate-free daughter cells. Interestingly, asymmetric segregation of protein aggregates seems to be correlated in many cases with an association of the aggregates with different cellular organelles ^8^. It has been suggested that an attachment to nucleus or vacuole, like in the cases of INQ and IPOD, resulting in a low number of relatively immobile aggregates, simply reduces the probability of inheritance to the new daughter cells ^30^. A recent report also described a prominent role of mitochondria in retention of cytosolic protein aggregates as tethering to mitochondria restricts their mobility and couples aggregate inheritance to the asymmetric distribution of damaged mitochondria ^31, 32^. The perinuclear association seems to serve this purpose in case of IMiQC.

In our system, the IMiQC represented a distinct form of functionless mitochondria, as judged by the absence of a membrane potential and their separation from the mitochondrial network. From our experiments it is clear that aggregate deposition and separation is highly dependent on mitochondrial dynamics, fission and fusion processes inside the mitochondrial network ^15^, because a deletion of yeast fission factor Fis1 led to cytotoxicity in presence of aggregation-prone polypeptides and a deletion of the fusion factor Fzo1 resulted in a higher number of aggregates. These observations support our conclusion that IMiQC formation requires a separation of the aggregates from the rest of the mitochondrial network combined with a gradual merger of the aggregate compartments into a single deposit site. However, it cannot be excluded that IMiQC formation involves the recently described generation of a specific form of mitochondrial vesicles ^33^, which potentially collects the aggregated polypeptides. An involvement of mitochondrial dynamics in protecting organelle quality has been already discussed in the context of neurodegenerative diseases that often comprise different instances of mitochondrial dysfunction. In mammalian cells, the cellular machinery controlling mitochondrial dynamics seems to allow a specific removal of damaged mitochondria while retaining intact organelles ^34^. Our results show that IMiQC formation is an important variant of general cyto-protective reactions involved in quality control of mitochondria. The importance of mitochondrial dysfunction has been increasingly recognized as a major factor in human pathologies, in particular neurodegeneration ^35^. After our identification of a specific mitochondrial aggregate sequestration mechanism that protects mitochondrial function and integrity under proteotoxic stress conditions, it will be of high interest to assess the relevance of IMiQC formation in a disease context.

## Acknowledgement

We thank Prof E. Hurt for providing the plasmid pRS314-NUP120-mCherry. We are grateful to Prof. E. Deuerling for the opportunity to perform the microscopic analysis in her laboratory. Work in the author’s laboratory was supported by the Deutsche Forschungsgemeinschaft (Grant VO 657/5-2 to W.V.).

## Author contributions

MB performed most of the experiments of this manuscript. CR performed the *in organello* translation experiment. GC performed the BN-PAGE experiments. WJ performed *in organello* degradation assays and thermotolerance experiments. AW and MS performed 2D-PAGE analysis of aggregate pellets. MB and WV designed the study, supervised the experiments and wrote the manuscript.

---

**Table I.**
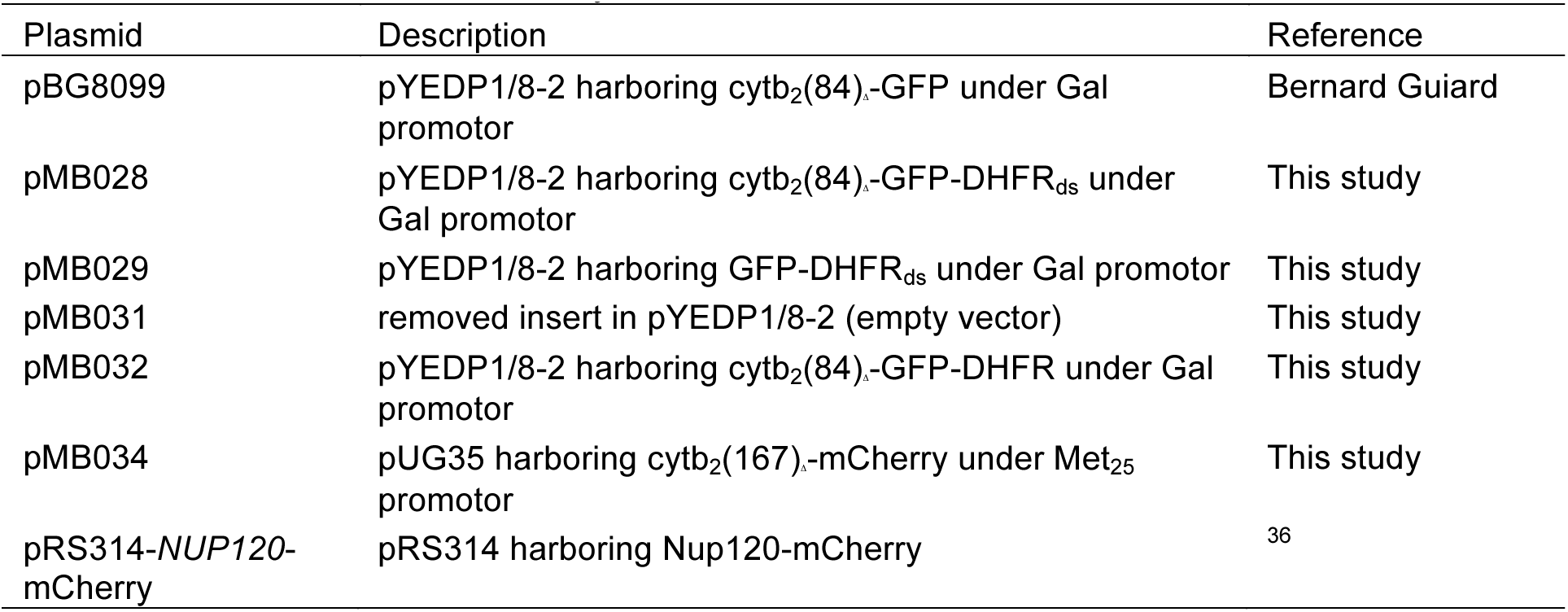
Plasmids used in this Study

**Table II.**
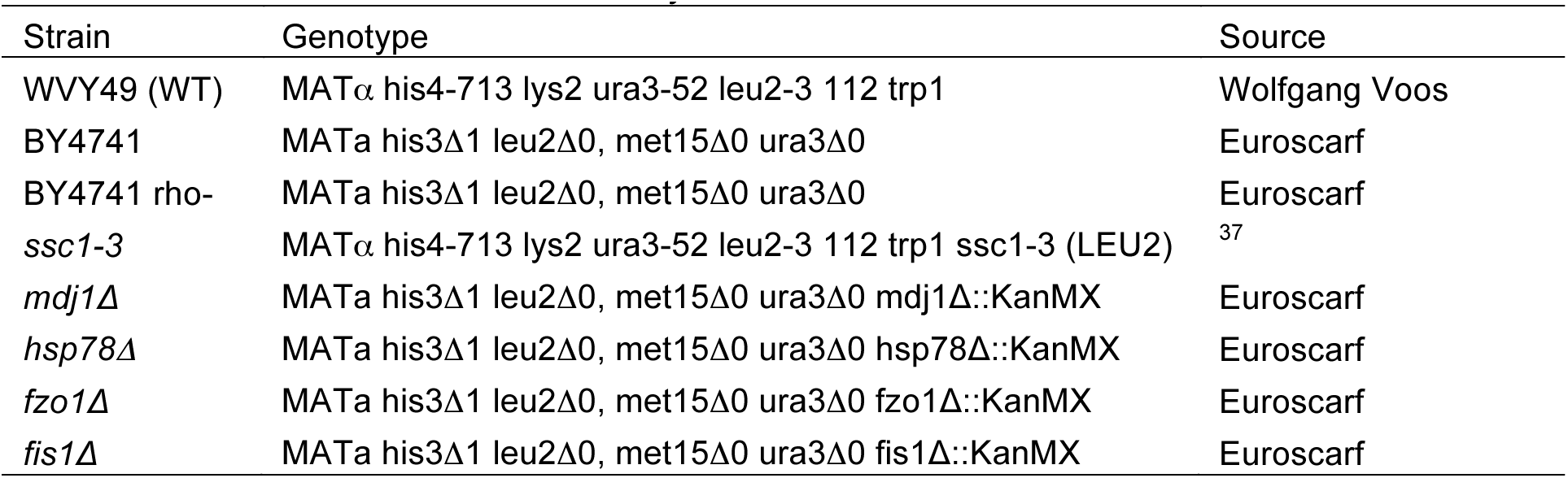
Yeast strains used in this Study

